# Mitochondrial dynamics resolve physical-social tensions and challenges through adaptable multi-objective optimisation

**DOI:** 10.64898/2026.09.22.753427

**Authors:** Joanna M. Chustecki, Iain G. Johnston

## Abstract

Complex biological behaviours often emerge when a system is faced with mutually incompatible priorities. In such situations, the field of multi-objective optimisation can provide an informative and predictive theoretical foundation for biology. Here we adopt this paradigm in exploring the rich dynamic behaviour of mitochondria inside cells. Using plant cells as a model system, we use physical modelling to characterise the “morphospace” of possible mitochondrial behaviours, and single-cell microscopy with video analysis and network modelling to characterise collective mitochondrial dynamics. We show that wildtype *Arabidopsis* mitochondrial dynamics near-optimally resolve a tradeoff between maintaining physical spacing and supporting biomolecular exchange. With existing and new experimental data, we show that these dynamics adapt under mutational and chemical challenges to support a rebalanced, but still near-optimal, resolution to this tradeoff under different densities of the mitochondrial population. We also show how an assumption of multi-objective optimisation supports inference of biological mechanisms before any data are observed, and discuss the potential of this multi-objective optimisation paradigm to form a broader theoretical framework of spatial cell biology.

## Introduction

Physical form and biological function are inextricably linked across scales in biology. As the “society” of cellular organelles is revealed in increasing detail by beautiful experiments (Guo et al., 2018; Perico & Sparkes, 2018; Valm et al., 2017; Viana et al., 2023; Wang & Mukherji, 2022), a theory of cellular organisation – and spatial cell biology – becomes a more feasible target (Chustecki & Johnston, 2024; Cohen et al., 2018; Jayashankar & Rafelski, 2014; Nurse, 2021; Wang & Mukherji, 2022). However, such a theory of “why” the cell is organised in a particular way requires study of the other ways it could possibly behave, to avoid traps of teleogical and proximal reasoning (Shellberg, 2001). In other words, the only comprehensive way to address the question “why does the cell do this” is by exploring “why doesn’t the cell do that”. Such a broad survey of possible alternatives and their utility is a challenging target with experimental approaches.

This in turn suggests a role for combining experiments with modelling approaches, where the consequences of putative behaviours can be explored free of implementational challenges, and possible mechanisms can be compared in the light of data (Kirk et al., 2013). Predictions from theories based on biological optimisation – argued to be “the only approach biology has for making predictions from first principles” (Sutherland, 2005) – can then be tested by asking whether observed biological behaviour performs better than a set of possible alternatives, with respect to some utility.

Here, we adopt this philosophy to explore a particular instance of rich, complex collective behaviour in cell biology: the dynamics of mitochondria inside plant cells. Plant mitochondria are essential for all plant (and hence animal) life, and exist largely as fragmented organelles exhibiting striking dynamic motion across the cytosol (Fig. 1) (Arimura et al., 2004; Arimura, 2018; Johnston, 2019; Logan, 2006, 2010; Logan & Leaver, 2000; I. Scott & Logan, 2011).

**Figure 1.**
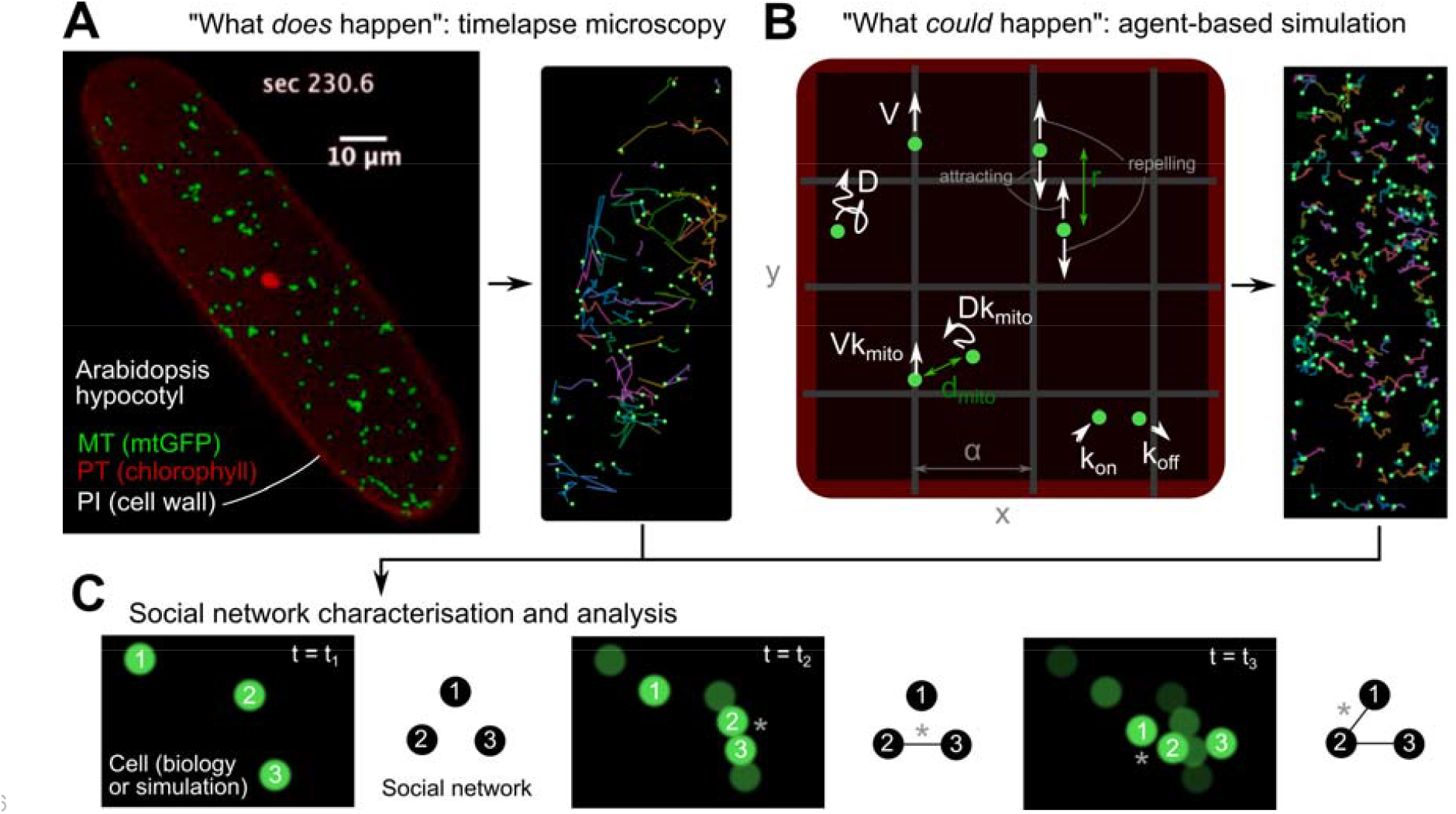
Model and analysis pipeline. **(A)** Cell biology: “what does happen”. Single-cell timelapse confocal microscopy in mtGFP *Arabidopsis* seedling hypocotyl followed by particle tracking identifies the trajectories of individual mitochondria. The imaging plane intersects the quasi-2D section of cytosol constrained between the cell wall and the vacuole. MT, mitochondrion; PT, plastid; PI, propidium iodide. **(B)** Physical simulation: “what could happen”. Mitochondrial motion is simulated, parameterised by a collection of physical processes including diffusion (constant D), ballistic motion on the cytoskeleton (speed V, association rate k_on_, dissociation rate k_off_), and between-mitochondrial influences (attraction or repulsion within distance r; scaling D and V by k_mito_ when mitochondria are within distance d_mito_). (see Methods). Simulation is scaled in space and time to reflect the cell biological samples. **(C)** Social network construction over time (t): a “social network” of mitochondrial encounters is constructed, where every node is a mitochondrion and an edge exists if two mitochondria have colocalised within a threshold distance (asterisks) over the recorded time course. “What does happen” vs “what could happen” can then be compared: summary statistics of mean minimum distance between mitochondria and mean degree in the social network are computed and compared across simulated and experimental observations. See also Supp. Videos 1-2 (<u>here</u> and <u>here</u>).

Perturbations to these dynamics are associated with severe plant phenotypes (Chustecki et al., 2022; El Zawily et al., 2014).

Previous work has explored optimisation in organelle dynamics, harnessing powerful approaches from physical and mathematical modelling (Hoitzing et al., 2015, 2019; Pain et al., 2019; Perico & Sparkes, 2018), with some exciting recent progress towards general principles (Amiri et al., 2023; Paul & Kollmannsberger, 2020). Many of these approaches consider a single cellular priority (for example, supporting spread through a network). In particular, the transport and spread of contents throughout mitochondrial structures of different topologies has been the subject of deep theoretical developments (Agrawal & Koslover, 2021; Chuphal et al., 2024; Holt et al., 2024, 2026; Z. C. Scott et al., 2021).

In previous work we implicitly proposed a contrasting picture of multiple objectives – where a system’s behaviour must resolve several competing priorities. Plant mitochondria contain only a subset of the mitochondrial genome and must therefore meet and exchange biomolecular content to acquire the machinery they need (Giannakis et al., 2022; Preuten et al., 2010). However, it is also beneficial for mitochondria to be evenly spaced through the cell, to support inter-organelle interactions and avoid heterogeneity of metabolites and other chemical species (Chustecki et al., 2021). Mitochondria cannot both be together (to exchange biomolecules) and apart (to preserve spacing) without moving, and our previous work showed that mitochondrial dynamics allow a resolution to these competing objectives (Chustecki et al., 2021).

But such a tradeoff could be resolved through many different modes of behaviour. The question of why plant mitochondria move the way they do therefore remains open. Connecting this multi-objective picture with an optimisation-based answer to “why” questions requires the paradigm of multi-objective optimisation – how a system can optimally resolve a tension between competing objectives (Collette & Siarry, 2004; Fieldsend & Everson, 2005). Here, we use this paradigm to explore the hypothesis that plant mitochondrial dynamics provide an adaptive, optimal resolution to these competing cellular tradeoffs.

## Results

### The morphospace of possible mitochondrial behaviours

Our workflow involves using single-cell microscopy to characterise the behaviour of real plant mitochondria (Fig. 1A), a physical model (Fig. 1B) to characterise the morphospace of possible mitochondrial behaviours, then quantifying the spacing and capacity for social exchange in all cases (Fig. 1C) to explore “what could happen” versus “what does happen”. To this end, we characterised the possible physico-social morphospace of plant mitochondria with over 2 × 10^5^ simulations of mitochondrial motion in model cells (Fig. 2A-D; Supp. Videos 1-2). We imposed observed physical constraints on the simulated behaviour, limiting diffusion and speed to within observed maxima from (Chustecki et al., 2021) and constraining cell geometry and mitochondrial density to observed ranges (see Methods). The resulting physically plausible morphospace displays a clear tradeoff between maintaining physical spacing and supporting encounters for biomolecular exchange (Fig. 3A). A Pareto-like front bounds the morphospace, giving the set of optimal resolutions to this tradeoff. The physical parameters that drive the system towards this Pareto front are maximum mitochondrial speed and propensity to move on the cytoskeleton (Supp. Fig. 1).

**Figure 2.**
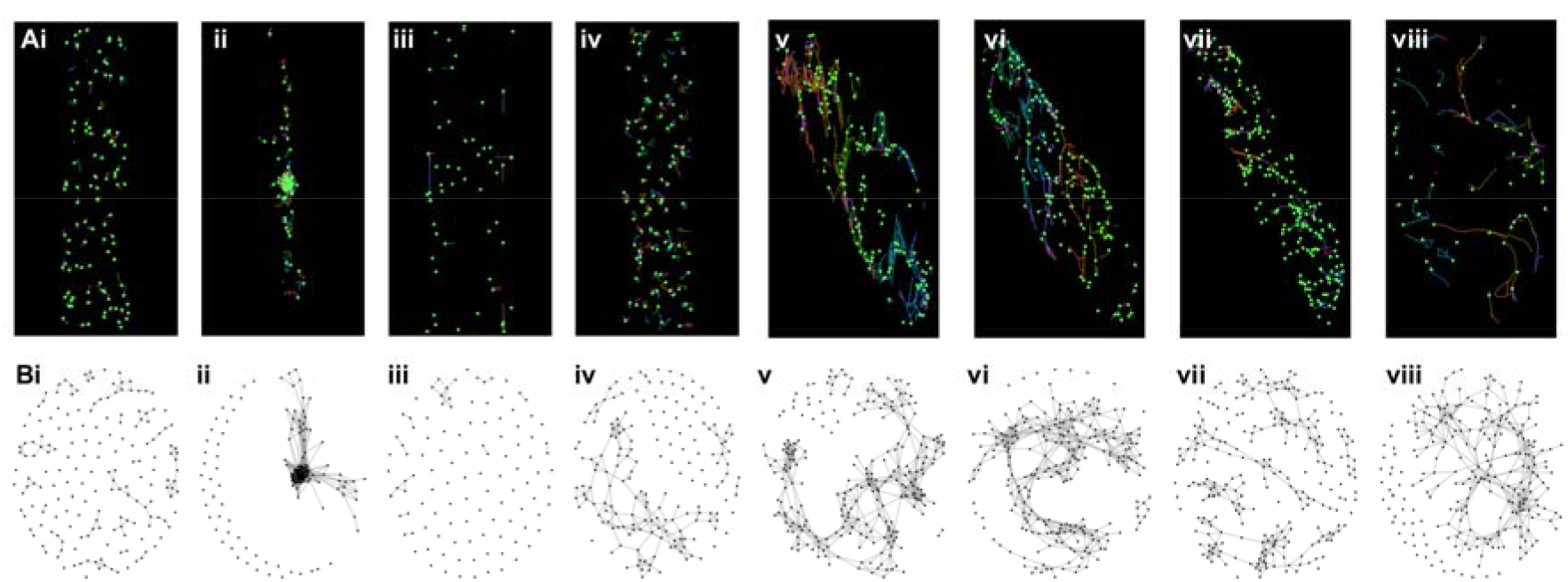
Simulation and observation of collective plant mitochondrial dynamics. **(A)** Mitochondrial dynamics and **(B)** resultant encounter networks. **(i-iv)** Example trajectories and social networks from different simulation parameterisations (“what could happen”), including parameterisation inducing (i) more static, (ii) more clustering, (iii) more cytoskeletal motion; (iv) more diffusion. **(v-viii)** Example trajectories and social networks from *Arabidopsis* microscopy (“what does happen”): (v) mtGFP (modelling wildtype); (vi) *msh1*; (vii) *friendly*; (viii) *msh1* treated with ciprofloxacin. The networks are dynamic objects that add connections over time; here, those in (i-iv) are measured after 100 simulation steps, and those in (v-viii) are compiled using the first 250 trajectories from each experiment. See also Supp. Videos 1-2 (<u>here</u> and <u>here</u>).

**Figure 3.**
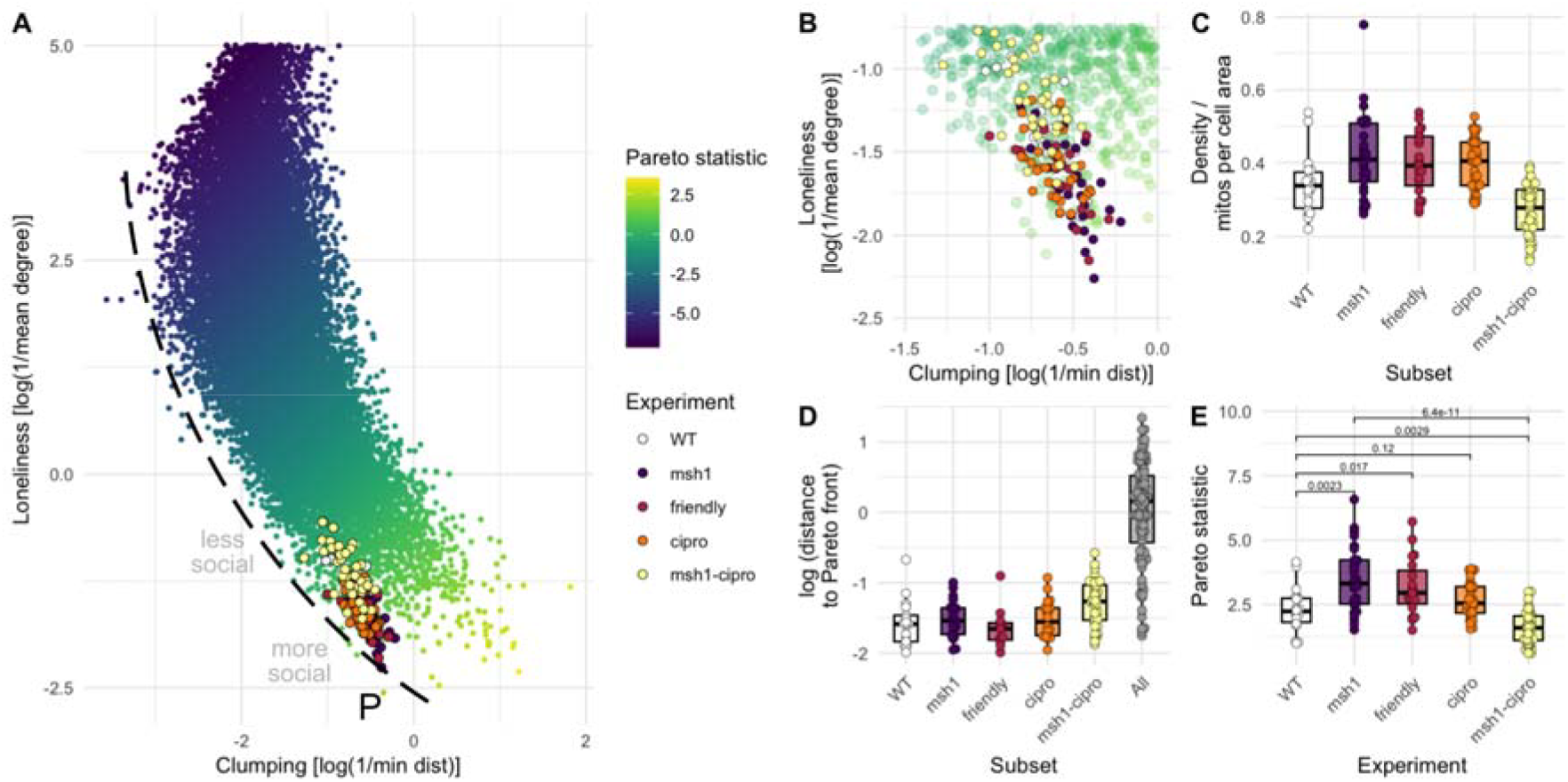
Near-optimal, adaptable resolution to a tradeoff between physical spacing and biomolecular exchange. **(A)** “What could happen” (physically plausible behaviour from simulation, blue-yellow points) vs “what does happen” (dynamics observed in experiments, points with borders) in the morphospace of mitochondrial priorities. Horizontal axis gives “clumping” statistic: low values indicate physically well-spread mitochondria. Vertical axis gives “loneliness” statistic: low values indicate well-connected encounter networks. Minimising both is impossible; the Pareto front (P) reflects the set of optimal tradeoffs, with the “Pareto statistic” characterising position along this front. Biological behaviour falls very near to the Pareto front. **(B)** Zoomed-in version of (A) showing proximity to the Pareto front and the shift in position between experimental lines. **(C)** Density of mitochondria in cells from different experiments. **(D)** Distance in morphospace to the Pareto front for each experimental observation and for all simulation parameterisations. Several experimental observations fall directly on P; distances for those that do not are shown. **(E)** Pareto statistic for experimental observations, showing the shift in the mutant lines and under chemical challenge.

Given that the physical model involves several sources of randomness (initial mitochondrial positions, random events, diffusive motion), each parameterisation produces a set of different points in the morphospace (Supp. Fig. 2). Computationally, this (finite) set corresponds to different choices of random seed; mathematically, the (infinite) set corresponds to different samples from the various distributions involved. The optimisation problem is then more strictly viewed as a set-valued optimisation problem (Khan et al., 2016) or multiobjective optimisation under uncertainty (Fieldsend & Everson, 2005) (see Discussion).

### Wildtype *Arabidopsis* mitochondria behaviour sits near the optimal Pareto front of a morphospace tradeoff

We next analysed the collective dynamics of mitochondria in *Arabidopsis* hypocotyl using the mtGFP line, providing a model for wildtype behaviour (Fig. 1A, 2E-H). We found that *Arabidopsis* mitochondria occupy a region of the morphospace almost exactly corresponding to the Pareto front (Fig. 3A-B), reflecting the optimal tradeoff (within physical limits) between spacing and exchange. In other words, over the range of physically possible collective behaviours that plant mitochondria could adopt, those observed in plant cells represent a near-optimal tradeoff between maintaining mitochondrial spacing and facilitating biomolecular exchange. The distances from experimental observations and the Pareto front are very small compared to the length scales in morphospace (median distance 0.19 units compared to spread around 2-4 units, Fig. 3B, D) and very different from distances from morphospace as a whole (median 6.0x closer to Pareto front, p < 2.2 × 10^-16^ from Mann-Whitney test).

Fig. 3 reflects an overview of the morphospace combined over different cell geometries and mitochondrial densities. Supp. Fig. 3 shows the subset of dynamic behaviours observed in the model when the parameterisation is restricted to match the geometry of each single-cell observation. Biological behaviour invariably falls around the Pareto front of each cell-specific subset of morphospace. Few instances of few parameterisations yield observations closer to optimality than the biologically observed behaviour: some examples are illustrated in Supp. Fig. 4.

### Mutant *Arabidopsis* mitochondria also occupy the optimal Pareto front but with shifted morphospace poise

We next analysed the collective dynamics of mitochondria in two *Arabidopsis* mutant lines, known to perturb mitochondrial population dynamics: the *friendly* mutant and the *msh1* mutant. A change in collective dynamics, favouring encounters by sacrificing physical spacing, has previously been reported in these mutants (Chustecki et al., 2021, 2022). Our new approach shows that these changes reflect a shift down the Pareto front (Fig. 3B-D; Supp. Fig. 3). The collective dynamics in the mutants still reflect a near-optimal tradeoff, but one rebalanced to favour exchange over spacing, commensurate with an increase of the density of the mitochondrial population in cells (Fig. 3C). For proximity to the Pareto front, no statistically detectable difference was observed between any of the three lines (wildtype, *msh1*, and *friendly*) (Fig. 3D); the Pareto statistic did not differ detectably between *msh1* and *friendly*, but both differed from the wildtype case (p = 0.0023 and 0.017 from Kruskal-Wallis with Mann-Whitney post hoc, Fig. 3E).

### Influence of chemical challenges on collective mitochondrial dynamics

Ciprofloxacin has been observed to compromise mtDNA integrity in plant cells, by inhibiting DNA gyrase causing random breaks (Schatz-Daas et al., 2022). To explore how such a chemical challenge might influence collective mitochondrial behaviour, we exposed both wildtype and *msh1* mutant plants to 0.5 µM ciprofloxacin on MS plates. The effect on wildtype was limited, inducing a small shift in the same direction as the *msh1* and *friendly* mutations (Fig. 3), but to a lesser extent which did not provide statistical support against the null hypothesis of no effect. However, ciprofloxacin treatment on *msh1* plants induced a strong effect in the opposite direction, where mitochondria reduced their encounters in favour of even spread (p = 6.4 × 10^-11^ from Mann-Whitney test, Fig. 3E), while remaining tightly constrained around the Pareto front (Fig. 3B, D). This likely reflects a change in cellular strategy under the combined genetic and chemical challenge: mitochondrial content is generally depleted, leading to lower cellular densities (Fig. 3C), and the collective dynamics adapt to support the optimal amount of interaction given this reduced density. Effectively, the line is forced upwards in the “isolated” direction in Fig. 3A, and physical arrangement is corresponding shifted leftwards to retain proximity to the Pareto front. These changes occurred without a dramatic change in the distribution of mitochondrial speeds (Supp. Fig. 5).

### Inferring mechanisms generating optimal, extreme, and biological behaviour

We next asked what parameterisations of our physical model gave behaviours most comparable to those observed in biology. For this process we used a rejection sampling implementation of Approximate Bayesian Computation (ABC) (Toni et al., 2008), reporting parameter sets that gave simulated behaviour whose statistics fall within a given threshold distance of biological observables (Methods, Fig. 4A). Several parameters are relatively unconstrained, meaning that a range of possible values are compatible with biologially observed behaviour. These include the distance at which position-based scaling of mitochondrial motion becomes active: as the highest posterior density falls in the region where this has no effect (k_mito_ = 1), the characteristic distance d_mito_ is thus quite unconstrained. Cytoskeletal attachment and detachment rates (k_on_ and k_off_) are individually quite freely varying, though unlikely to be zero; their ratio k_on_ / k_off_ is more tightly constrained around 1, suggesting a mixture of cytoskeletal and diffusive motion. Physical features of the cell are also constrained, with dimensions, mitochondrial density, and cytoskeletal spacing all having pronounced posterior peaks. The diffusion constant distribution has a peak at 0.1 µm^2^ s^-1^, and the characteristic cytoskeletal speed has a peak at 1 µm s^-1^, compatible with observations from previous studies (Chustecki et al., 2021; Zheng et al., 2009). This suggests that cytoskeletal structure, rapid cytoskeletal motion, and limited but nonzero diffusion are important for resolving the spacing-social tradeoff we identify.

**Figure 4.**
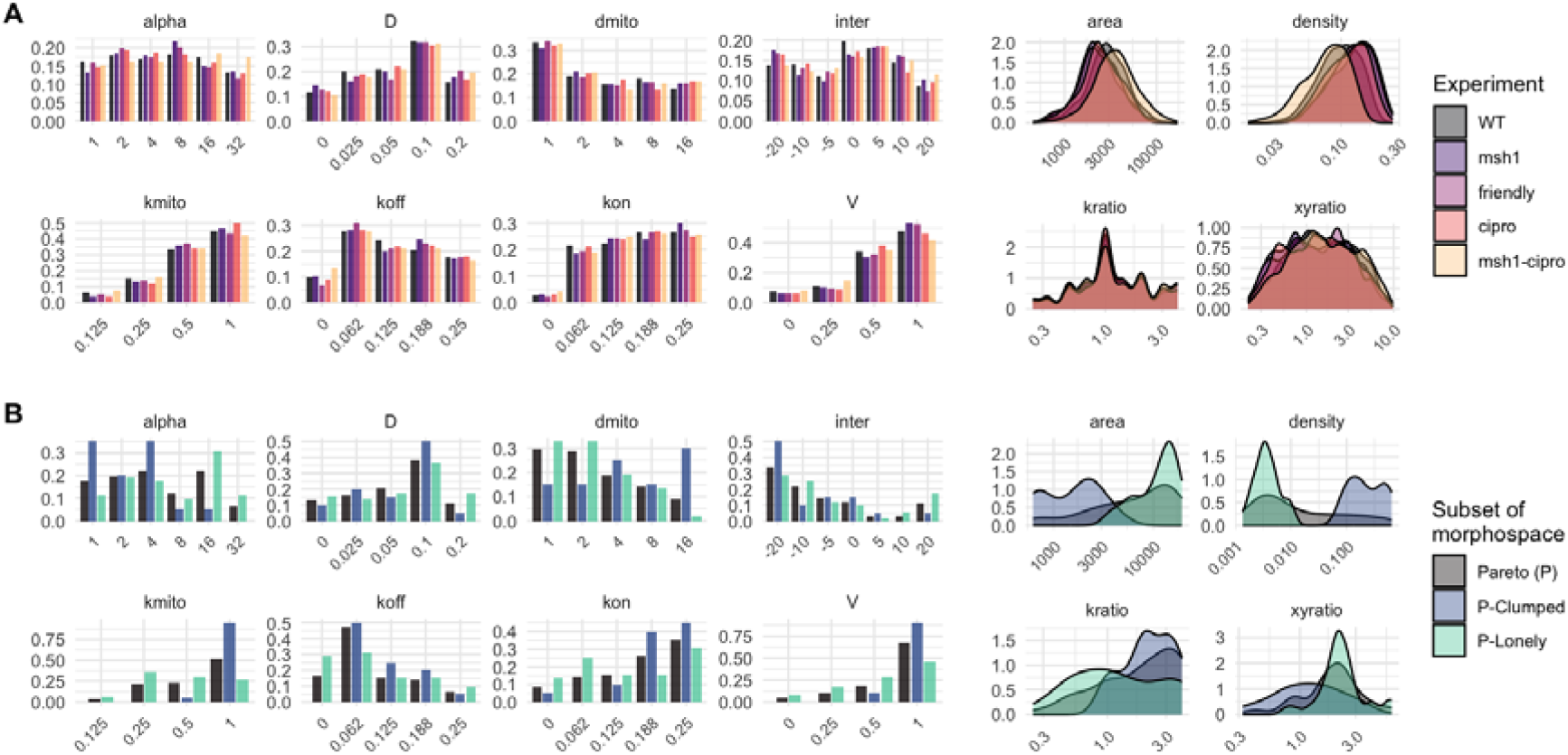
Inference of generating parameters with and without data. Posterior distributions from approximate Bayesian computation on the physical parameters in the simulation model. Parameters are diffusion constant D [μm s^-1^]; cytoskeletal spacing α [μm]; trafficking speed V [μm s^-1^]; cytoskeletal binding and unbinding rates k_on_ and k_off_ [s^-1^]; interference distance d_mito_ [μm]; interference speed scaling k_mito_; interaction radius r [μm] and interaction polarity. “inter” is the product of interaction radius r and polarity of interaction (negative values for attraction, positive for repulsion); “area”, “xyratio”, and “density” are respectively cell area [μm^2^], aspect ratio, and mitochondrial density [μm^-2^]; “kratio” is k_on_ / k_off_. **(A)** Inference using biological observations. Different colours give different plant lines. **(B)** Inference without data. “Pareto (P)” gives posteriors P_ε_(θ|P) resulting solely from the assumption that biology occupies the Pareto front P, without seeing any specific observations. “P-Clumped” gives the subset of parameterisations on the Pareto front that produce clumping > -1; “P-Lonely” gives the subset on the Pareto front that produce clumping < -2.1 (see Fig. 3A).

Pursuing the multi-objective optimisation picture, we next considered a more unusual question. It is clear from Fig. 3A that the region around the Pareto front, reflecting an optimal tradeoff between spacing and exchange, constitutes a small region of morphospace overall. It could follow that only a restricted set of mechanistic parameters would yield near-optimal behaviours. We therefore next asked to what extent we could estimate parameters conditioned not on observation, but solely on the insight that a system likely adopts a near-optimal resolution to competing priorities.

For convenience we pursue the ABC picture. In standard ABC, as above, a likelihood calculation is replaced by comparing data D with pseudodata D(θ) simulated from the model with parameters θ. If a distance between the pseudodata and the real data is under a threshold ε, we accept θ as a sample from the “posterior”: P_ε_(θ|D) ∼ I(|D – D(θ)| < ε) P(θ), where I is an indicator function (Beaumont et al., 2002; Toni et al., 2008). But now consider a situation where we have no specific data D but believe that such data would lie within a set P (here the Pareto front, or some region around it). We could then consider using P_ε_(θ|P) ∼ I(min_p_∈P |p – D(θ)| < ε) P(θ). In other words, accepting parameterisation θ as a sample from the posterior if it generates pseudodata that falls within ε of our target set.

Sampling the parameters corresponding exactly to the elements of the Pareto front in Fig. 3A yield “posteriors” P_ε=0_(θ|P) that already constrain some parameter estimates for θ (Fig. 4B). In particular, this “inference without data” suggests a sparse cytoskeleton, intermediate diffusion constant, nonzero cytoskeletal attachment and detachment, and high cytoskeletal speed to be requisites for optimal behaviour, all of which are consistent with the inference with data in Fig. 4A. Other inferred features of the Pareto set are less informative, because of the range of “strategies” by which a parameterisations can occupy the Pareto front (Supp. Fig. 4). A range of interaction scales and directions, and cell densities, for example, correspond to either “isolated” or “clumped” regions of the Pareto front.

Following this, we asked which parameter values were particularly associated with extreme ends of the Pareto front (sacrificing spacing for interaction, or vice versa). We observe, intuively, that “lonely” behaviour corresponds to large cells with sparse mitochondrial populations, higher cytoskeletal detachment than attachment, low cytoskeletal speeds and more repulsion between mitochondria. Conversely, more “clumped” behaviour corresponds to high cytoskeletal attachment and speed, diffusion rates, and higher densities. These trends agree with the general position of biological observations towards the “clumped” regime, with the msh1 and friendly lines displaying slightly more “clumping” parameter characteristics, and msh1-cipro displaying slightly more “lonely” characteristics (Fig. 4A).

Hence, a “data-free inference” step based only on the notion of Pareto optimality already constrains several key parameters of the dynamic system to a reduced, biologically consistent region of space.

## Discussion

We have shown that, relative to the morphospace of physically plausible mitochondrial dynamics, wildtype, mutant, and chemically challenged cells provide near-optimal resolutions to a tradeoff between physical spacing and biomolecular exchange. This does not, of course, mean that this is the complete set of priorities that drive the evolution of mitochondrial dynamic control. Mitochondrial ultrastructure fulfils many other cellular functions, several in opposition, and varying by cell type, environment, and more features (Chustecki & Johnston, 2024). Our goal is not to claim a fully causal theory of mitochondrial dynamics, but to demonstrate an optimal resolution to one of these tensions.

What predictions can be made from this theory? Our results suggest that cells can adapt their collective mitochondrial dynamics in response to different challenges. Fusion mutants, for example, will struggle to exchange contents and are thus predicted to occupy a region on the Pareto front in Fig. 3A shifted towards clumping – sacrificing spacing to promote interactions and attempt to maintain exchange between mitochondria. This is supported by increased mitochondrial density in *miro2* mutants, large clusters with “megamitochondria” in PMF overexpression lines, and large groups of individual mitochondria in *fmt* mutants (El Zawily et al., 2014; Kenneally et al., 2026; White et al., 2020). In situations where sharing of contents is critical, like the shoot apical meristem, our theory would again predict that cells occupy a position on the Pareto front sacrificing spacing, compatible with the observation of highly fused structures in that tissue (Seguí-Simarro & Staehelin, 2009). Observations of collective dynamics far from the Pareto front would challenge our picture, suggesting that dynamics are responding to other cellular priorities (or none).

More broadly, our approach predicts that the resolution of physical-social tensions may be one important goal of the complex and incompletely understood mitochondrial dynamics observed in plant cells, and more broadly across species (Hoitzing et al., 2015). Biomolecular exchange through social interaction has several adaptive advantages, including complementation and genome maintenance (Arimura, 2018; Giannakis et al., 2022). This is in agreement, for example, with our and previous *msh1* observations: the mutation challenges mtDNA integrity, and the physical response we observe is compatible with a response promoting exchange (Chustecki et al., 2022). Our prediction that the maintenance of physical spacing is a competing priority comes from spatial considerations of uniform ATP supply (Kajita et al., 2024; Kumar & Johnston, 2026), avoidance of local reactive oxygen species buildup, and facilitation of inter-organelle communication (Chustecki & Johnston, 2024; Cohen et al., 2018). Interestingly, experimentally increased or arrested mitochondrial movement impacts their functional integrity and the motion of other organelles, demonstrating that even spacing and positioning is tightly regulated, predictable and important for morphological homeostasis (Gu et al., 2026).

Our approach has largely treated cellular (and simulation) noise as a statistical inconvenience rather than an intrinsic feature of the systems involved. As mentioned previously, the stochastic nature of these behaviour would suggest a further role for set-valued optimisation and multi-objective optimisation under uncertainty (Fieldsend & Everson, 2005; Khan et al., 2016) (Supp. Fig. 2). Stochastic optimal control (Wiegerinck et al., 2006) – previously applied to mitochondrial populations (Hoitzing et al., 2019) – also has potential to explore the optimal control of such systems in the face of inevitable cellular noise.

The characterisation of the morphospace of possible behaviours is dependent on our choice of model for mitochondrial dynamics. If, for example, we allow mitochondrial speed to be unconstrained by physical observations, better resolutions to the physical-social tension (simultaneously lower isolation and clumping in Fig. 3A) are readily achievable. We have attempted to be as general as possible while respectively physical reality and retaining tractability: our model supports the presence (over varying magnitudes) or absence of multiple different physical mechanisms in the cell, with flexible cell geometries and mitochondrial populations, informed by previous studies on plant mitochondrial dynamics (Arimura, 2018; Chustecki et al., 2021; I. Scott & Logan, 2011). However, it is possible that a model considering other mechanisms would yield a different morphospace structure; a model selection perspective specifically considering the microscopic processes involved would be a valuable future direction for this research (Chustecki & Johnston, 2024; Kirk et al., 2013).

More generally, we hope to highlight the power and applicability of multi-objective optimisation and other tools from operations research (OR) in cell biology (Hafner & Rieger, 2018; Helbing et al., 2009; Li & Zhang, 2022; Park et al., 2023). OR in the human world (with cellular metaphors aligned with this research in brackets) addresses what resource to invest in power plants (bioenergetic organelles), factories (ribosomes), a transport network (the cytoskeleton), communications (signalling pathways), facilitating trade through infrastructure (cytoskeleton transport of organelles), avoiding pollution with decentralization (spreading active organelles), zoning to control the proportion of different functions, and so on.

Feedback processes must exist to respond to changes in environment and economic supply and demand. OR exploits control theory, maths, and simulation to design optimal and robust structures in these contexts. The biological cell, faced with similar priorities of resource allocation, robustness, and responsiveness, may well have evolved control mechanisms that share comparable characteristics. As more data on spatial biology becomes available, an “optimal planning” theory of the society of cellular substructure may be a feasible and fruitful avenue of research (Chustecki & Johnston, 2024; Valm et al., 2017; Wang & Mukherji, 2022).

## Methods

### Physical simulation

We take the physical model from (Chustecki et al., 2021) as a foundation (Fig. 1A). N mitochondria exist in a rectangular 2D cell of dimensions X × Y [μm]. Mitochondria diffuse with coefficient D [μm^2^ s^-1^]. The cytoskeleton is modelled as a collection of strands that run horizontally and vertically throughout the cell, regularly spaced by distance α [μm]; an unattached mitochondrion within 3 μm of a strand can with rate k_on_ [s^-1^], and an attached mitochondrion detaches with rate k_off_ [s^-1^]. Attached mitochondria move unidirectionally and ballistically along the cytoskeleton with speed V [μm s^-1^]. If within a threshold distance d_mito_ [μm] of another mitochondrion, mitochondrial speed is scaled by a factor k_mito_. For generalising beyond the cases in (Chustecki et al., 2021), we (following recent modelling work (Kajita et al., 2024; Kumar & Johnston, 2026)) introduce an interaction term between mitochondria, which acts either to attract (parameter value -1) or repel (parameter value 1) a mitochondrion to or from the centre of mass of mitochondria within a threshold distance r [μm]. To model observed movement of mitochondria in and out of the imaging plane in microscopy (Chustecki et al., 2021), we allow mitochondria to transition into a “detectable” state and out to an “undetectable” state, with rates ρ_in_ and ρ_out_ = 0.01 s^-1^ respectively, with a mean 90% of detectable mitochondria, to match microscopy observations. We also vary cell geometry by drawing dimensions from distributions X ∼ U(0, 200) μm, Y ∼ U(0, 100) μm, and N ∼ U(0, 200) – capturing the range of values observed in source data (Chustecki et al., 2021, 2022), and employ reflecting boundary conditions inside the model cell. Mitochondria are randomly arranged throughout the cell at t = 0 s and simulated for 200 s equilibration “burn-in” then 200 s of observed behaviour, to match the video data (see below).

For this analysis, we systematically span parameter space over the following ranges. D ∈ [0, 0.1] μm^2^ s^-1^ (from observation of maximum ∼0.15 μm^2^ s^-1^ in (Chustecki et al., 2021)); α ∈ [1, 16] μm (reflecting a spectrum from a dense network to few strands per cell); V ∈ [0.025, 1] μm s^-1^ (from observation of maximum ∼1 μm s^-1^ in (Chustecki et al., 2021)); k_on_ ∈ [0, 0.25] s^-1^; k_off_ ∈ [0, 0.25] s^-1^; d_mito_ ∈ [1, 16] μm, k_mito_ ∈ [0.125, 1], r ∈ [0, 20] μm (for both attractive and repulsive interactions). To enforce physical limits on mitochondrial speed, we impose an universal speed limit of 0.2 μm s^-1^.

### Plant culture

Seeds of Col-0 and *friendly* mutant *Arabidopsis thaliana* with mitochondrial-targeted GFP were kindly provided by Prof. David Logan (El Zawily et al., 2014; Logan & Leaver, 2000), and used with mtGFP-*msh1* seeds from (Chustecki et al., 2022). Seeds were surface sterilized in 50% (v/v) household bleach solution for 4 minutes with continual inversion, rinsed three times with sterile water, and plated onto ½ MS Agar. Plated seeds were stratified in the dark for 2 days at 4°C. Seedlings were grown in 16hr light/8hr dark at 21°C for 4-5 days before use.

### Confocal microscopy and video analysis

After 4–5 d, seedlings were taken for imaging and, prior to mounting, stained with 10 µM propidium iodide (PI) solution for 3 min to capture the cell wall. Simple mounting of whole seedlings on microscope slides with coverslips was used (modified from (Ekanayake et al., 2015)). In order to minimize the effects of hypoxia and physical stress on the seedling, imaging was undertaken in <10 min after the coverslip was added.

We used two microscopes for image capture, the inverted Zeiss 710 laser scanning confocal (mtGFP, mtGFP-*friendly*, mtGFP-*msh1*), and the upright Nikon A1-NiE confocal (mtGFP, mtGFP-*msh1*, ciprofloxacin treatments). Images were captured in a similar manner. On the Zeiss microscope, channels used were: GFP, Ex: 488nm, Em: 535.5nm (494-578nm), Chlorophyll and PI, Ex: 543nm, Em: 679.5 nm (578–718 nm). On the Nikon channels used were: GFP, Ex: 488□ nm, Em: 525 (500–550), PI, Ex: 561, Em: 595 (570–620), Chlorophyll, Ex: 640□nm, Em: 700 (663–738). For all analysis, only GFP channels were used to quantify mitochondrial location and motility. For Zeiss images, resolution was 5 pixel/µm, for Nikon, 4.83 pixels/µm. Time lapses were taken, with intervals of 1.93s for the Zeiss system, and 2.21s for the Nikon system. These time intervals and length scales have been taken into account for all downstream analysis.

For image analysis, single cells were cropped using the PI cell wall outline with Fiji (Schindelin et al., 2012). To counter the occasional sample drift within time-lapse videos, drift correction was applied with default settings, using the cell outline via the PI channel as the stability landmark (Correct 3D drift, FIJI, ImageJ 2.1.0; (Parslow et al., 2014)).

Following (Chustecki et al., 2021), tracking of individual mitochondria was done using Trackmate (Tinevez et al., 2017) in ImageJ 2.0.0. The LoG detector was used, with typical settings being 1 µm blob diameters (the typical size of a mitochondrion), although 0.8 µm was occasionally used for lower signal samples, or 1.1µm for higher. The detection threshold was set between 1.5 and 8, and filters were applied on spots if necessary. The Simple LAP Tracker was run with a linking max distance of 4 μm (3 µm used for a few samples), gap-closing distance of 5 μm (4 µm used for a few samples), and gap-closing max frame gap of two frames. For each sample, the quality of overlaying detection for mitochondria was scrutinized, and occasional tracks were edited for precision.

### Ciprofloxacin treatment

Seedlings were grown on ½ strength MS plates (pH 5.5) with 0.5µM Ciprofloxacin (Combi-Blocks, #QE-1554). Ciprofloxacin stock powder was dissolved in MES Hydrate (pH 4.5), and diluted to a working stock of 0.5mM. Growth conditions were as above, stratified in the dark for 2 days at 4°C, and grown in 16hr light/8hr dark at 21°C for 4-5 days before use.

### Trajectory and social network analysis

Both simulations and videos were analysed over 200 s, with a sampling interval of 2 s (one frame in confocal microscopy, one sampling event in the simulation). Mitochondrial colocalisations within a threshold distance of 1.6 μm were recorded as encounters, corresponding to an edge in the encounter network. At each sampled snapshot, we recorded the minimum distance between each mitochondrion and its nearest neighbour and averaged this over mitochondria and snapshots to get the mean minimum distance for a cell (observed or simulated). We took the mean degree of nodes in the encounter network established over the 200s interval as the mean degree for a cell. To express these quantities in a Pareto framework, we define “clumping” = log(1 / mean minimum distance) and “loneliness” = log(1 / mean degree) as quantities that our hypothesised tradeoff should minimise. Clumping is the inverse of (desirable) physical spacing, and loneliness is the inverse of (desirable) social exchange capacity.

### Approximate Bayesian computation

We use the summary statistic |D_meanmin_ – D_meanmin_(θ)| + |D_meandegree_ - D_meandegree_(θ)| to compare real-world data D with pseudodata D(θ). The subscripts denote the particular observations of mean minimum distance and mean degree, aligning with the physical-social focus of our research. We use ε = 0.1 as a threshold for accepting a sample, following empirical observation balancing goodness of fit with reasonable probability of generating accepted samples.

## Data and code availability

All data from the ciprofloxacin experiments, and code for the analysis is available at https://github.com/StochasticBiology/optimal-mitos. Data from other experiments, and the pipeline for social network extraction, is at https://github.com/StochasticBiology/plant-mito-dynamics (Chustecki et al., 2022).

## Acknowledgements

The authors gratefully acknowledge support from and discussions with colleagues including Alan Christensen, Nick Jones, David Logan, and Markus Schwarzlander. This project has received funding from the European Research Council (ERC) under the European Union’s Horizon 2020 Research and Innovation Programme [Grant agreement No. 805046 (EvoConBiO) to I.G.J.]. This work was supported by the Research Council of Norway through the FRIPRO program [project numbers 357812 and 360449 (MitoPhyto) to I.G.J.]. This work was also supported by a WiRE Fellowship at the University of Münster to J.M.C. This work was also supported by a grant from the National Science Foundation to Professor Alan Christensen (MCB-1933590). The UNL Microscopy Core Facility is supported by federal funding from the NIH COBRE program (P20 GM113126). Major support was from a University of Nebraska Foundation fund in memory of Frank and Edith Christensen (01146140, to Professor Alan Christensen). I.G.J. thanks the Rudolf Peierls Centre for Theoretical Physics at the University of Oxford for hosting a sabbatical visit during work on this project, and L. Meltzers Universitetsstiftelse from the University of Bergen for supporting this visit.

## Supplementary Information

**Supplementary Figure 1.**
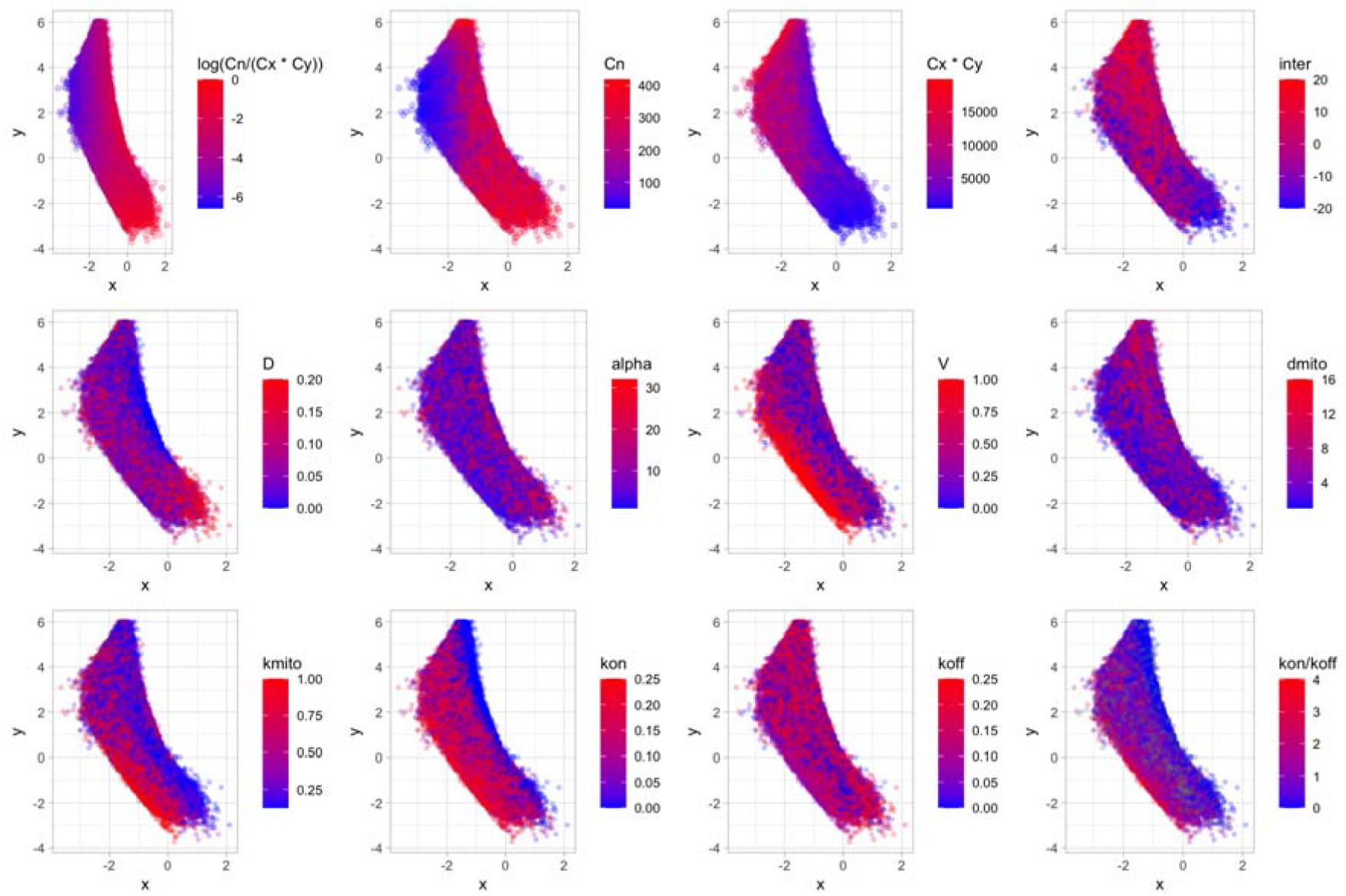
Physical-social morphospace with different simulation parameters. In each plot, every point is a simulation output in the clumping-loneliness morphospace from Fig. 3. The value of the given physical parameter for each simulation is plotted as the colour scale. Mitochondrial density, for example, in part determines the position in the clumping-loneliness tradeoff. Higher ballistic velocity V tends to drive the system towards the optimal Pareto front.

**Supplementary Figure 2.**
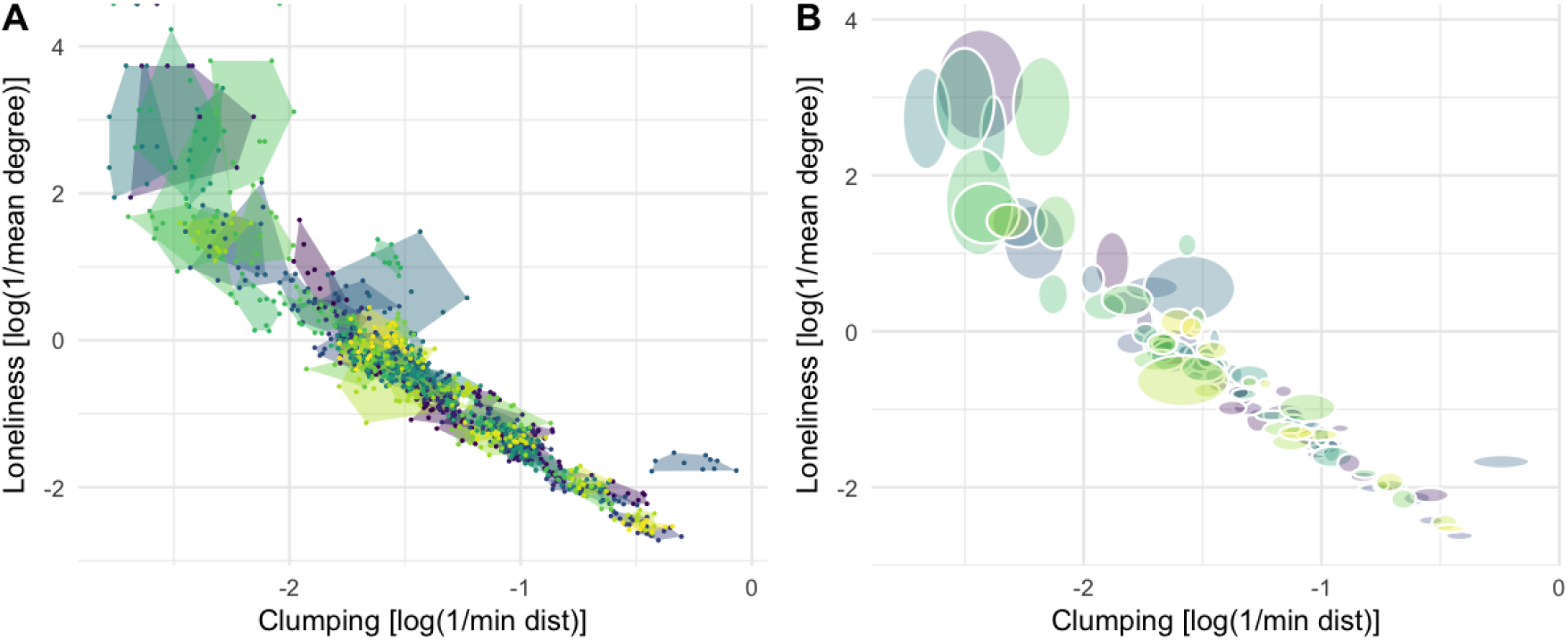
Set-valued optimisation. As each simulation is stochastic (for example, with random initial conditions and diffusive steps), each parameterisation of the model generates a set of points in physical-social morphospace. Here two visualisations of this set-valued behaviour are given for 10 samples from each of a subset of 100 parameterisations: **(A)** convex hulls of the set arising from each parameterisation, and **(B)** ellipses centred on the mean and with axes giving the standard error on the mean behaviour.

**Supplementary Figure 3.**
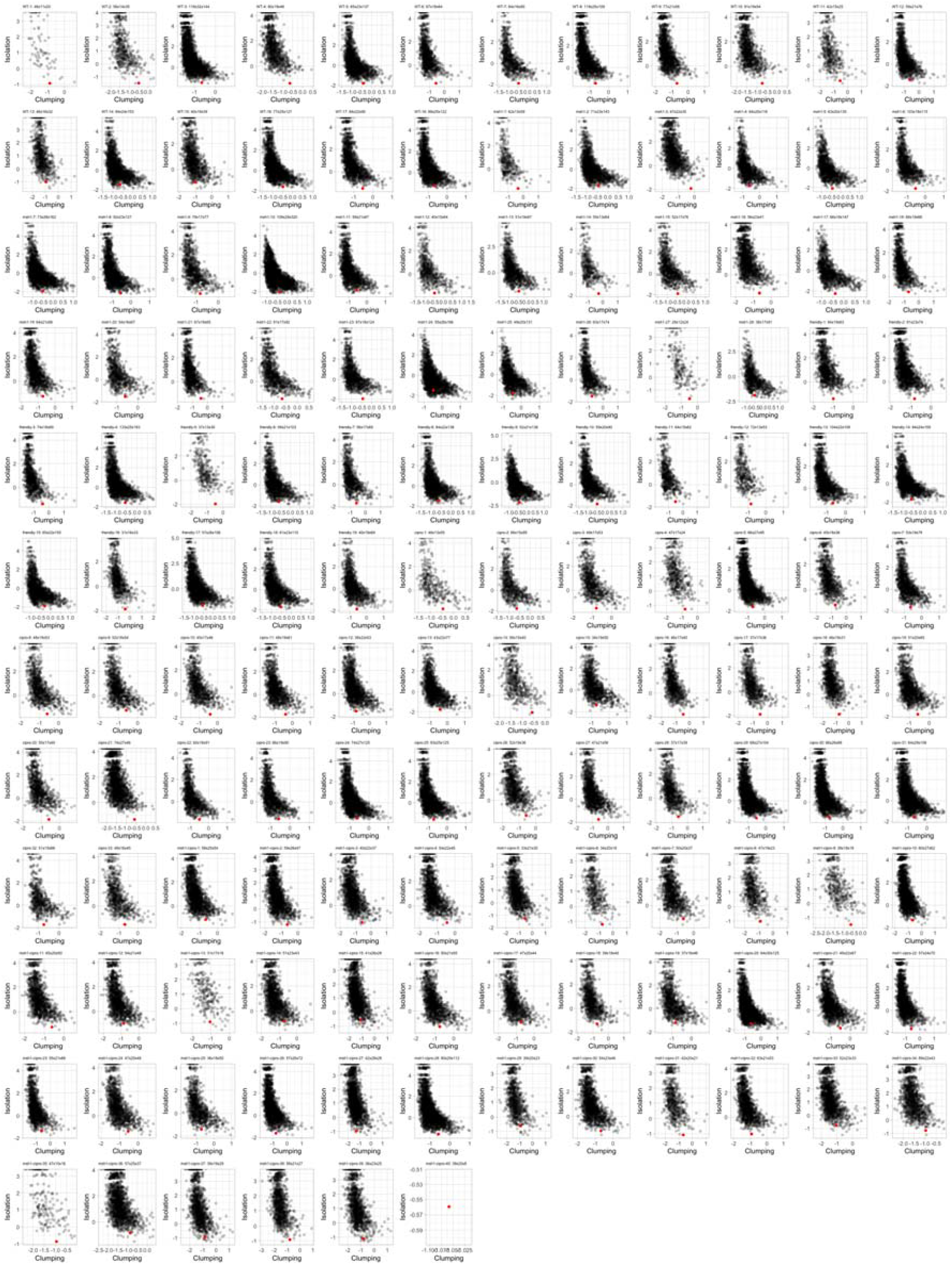
Cell-geometry-specific morphospaces. Each panel corresponds to one single-cell experimental observation. The subset of simulation parameterisations matching that cell’s geometry (cell dimensions within 10um, mitochondrial count within 10) are visualised in the physical-social morphospace (grey) and the experiment’s position in morphospace is given in red.

**Supplementary Figure 4.**
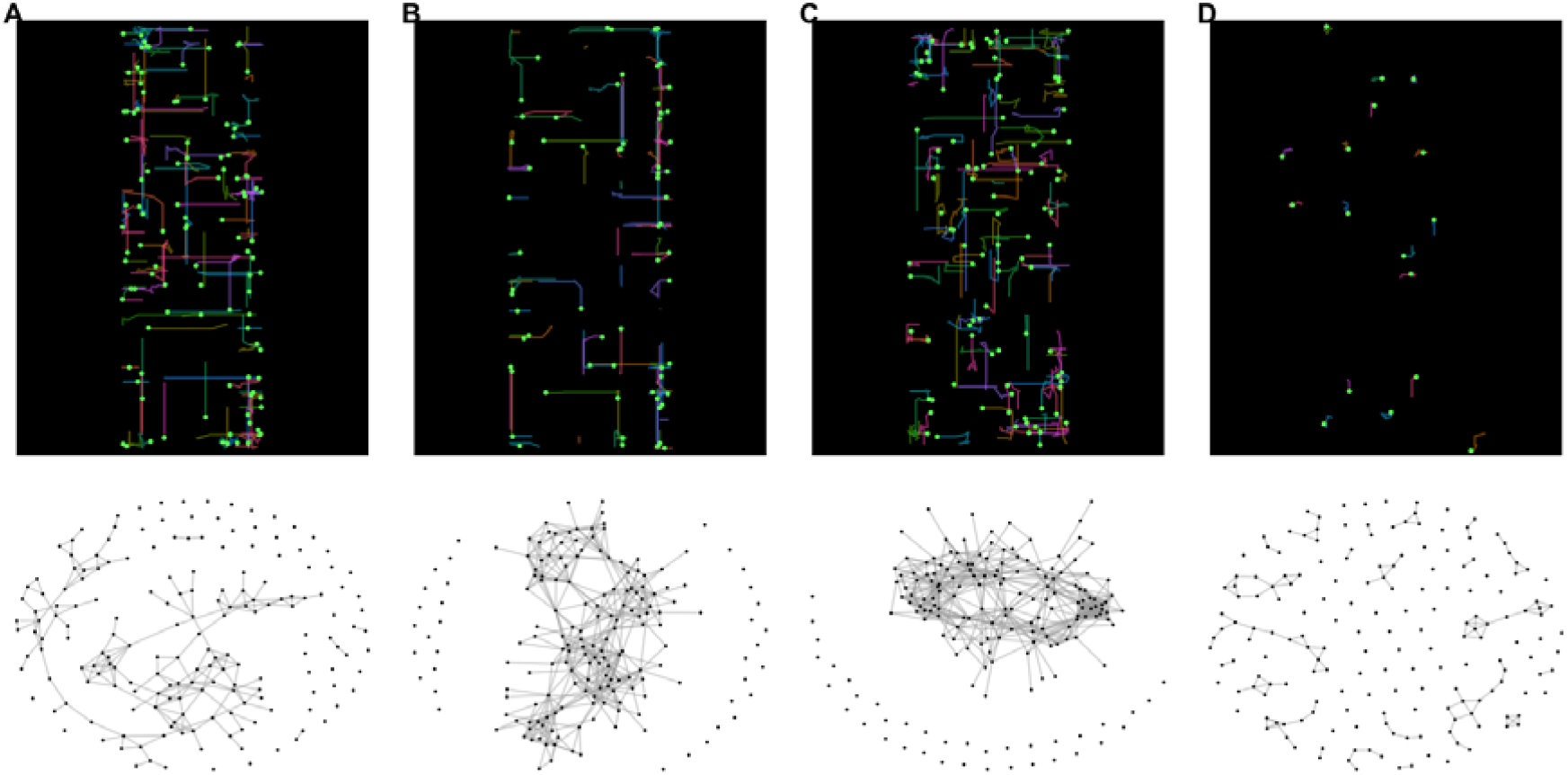
Illustration of simulations on the Pareto front. **(A-B)** Two simulations from parameterisations on the Pareto front (Fig. 3A) near the region occupied by biological observations. **(C)** Simulation from parameterisation on the Pareto front in the highly social, but clumped regime. **(D)** Simulation from parameterisation on the Pareto front in the highly-spread, but isolated regime.

**Supplementary Figure 5.**
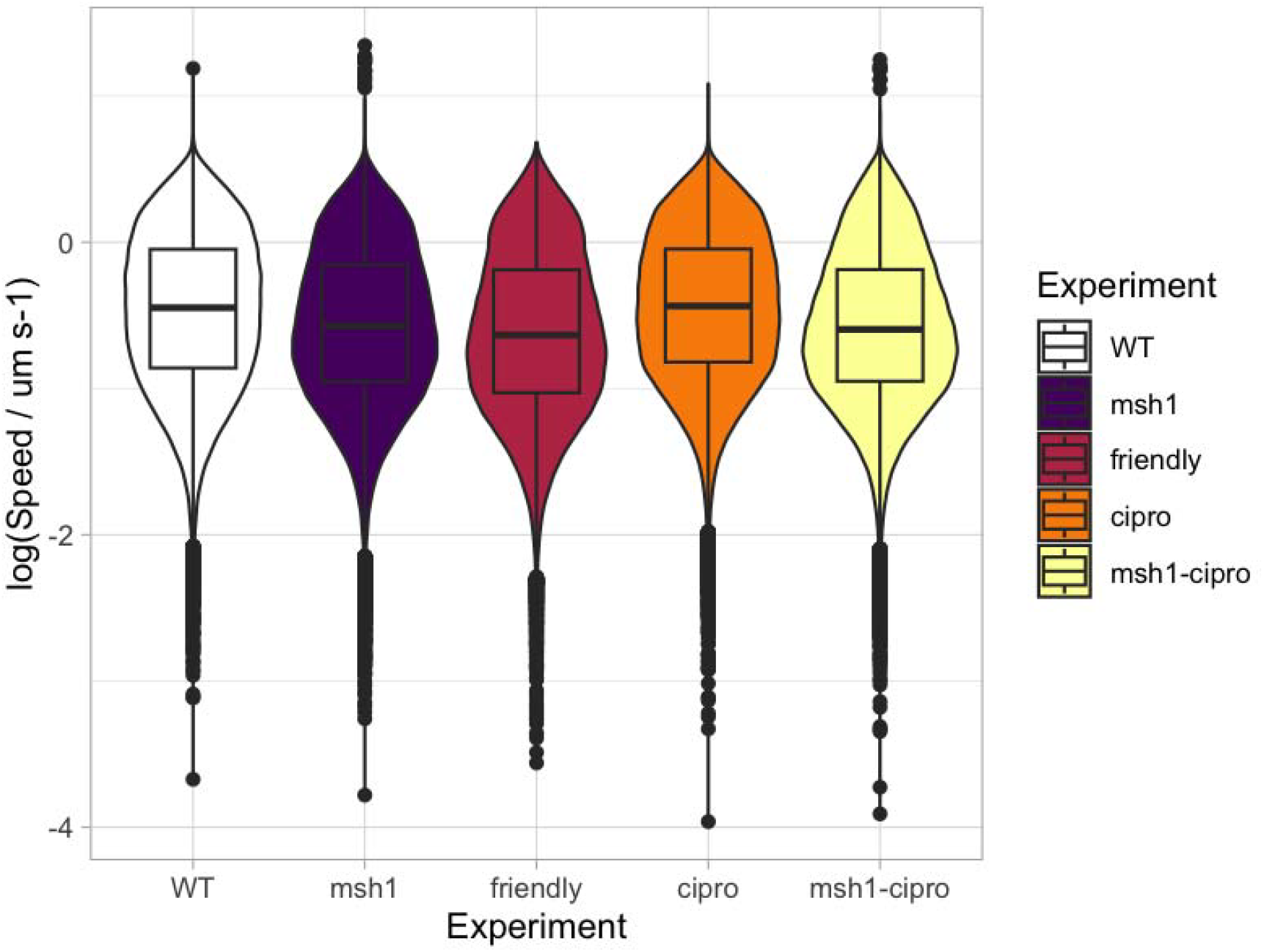
Distributions of mitochondrial speeds across the different experiments.

**Supplementary Video 1.**
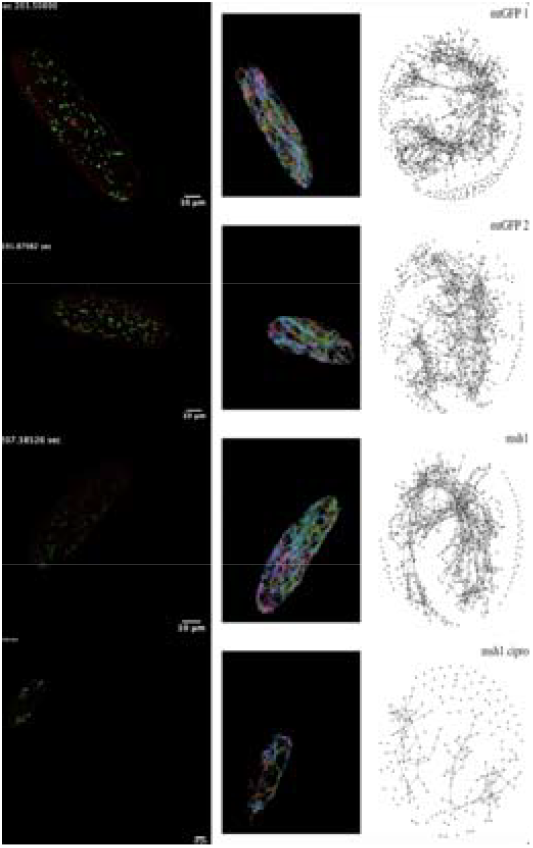
Example videos of observed mitochondrial dynamics and social network construction. Videos of the raw microscopy imaging, the segmented and tracked particles from video analysis, and the consequent social network evolving over time. From top to bottom: two mtGFP cells (modelling wildtype), *msh1* mutant, *msh1-cipro* treatment. https://github.com/StochasticBiology/optimal-mitos/blob/main/summary-videos/several-experiments.mp4

**Supplementary Video 2.**
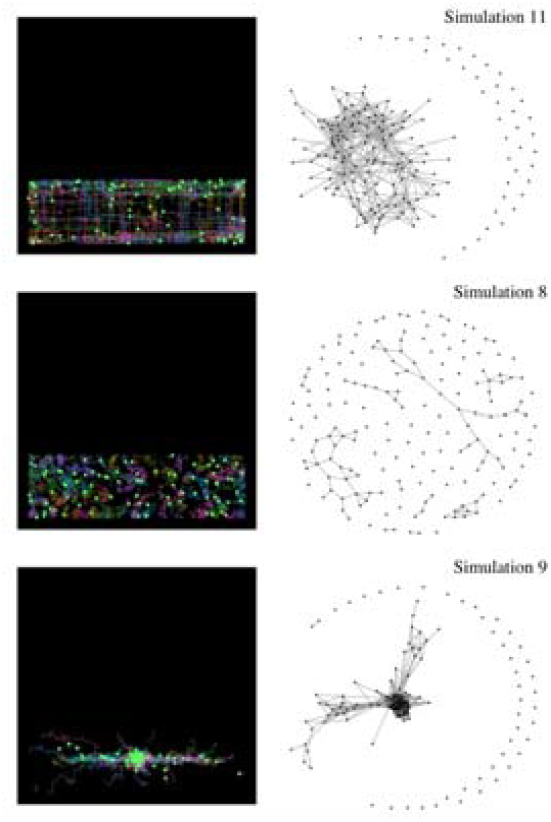
Example video of simulated mitochondrial dynamics and social network construction. (top) A parameterisation with a combination of cytoskeletal motion and diffusion, giving moderate spread and social connectivity. (centre) A parameterisation favouring physical spread over social connectivity. (bottom) A parameterisation favouring social connectivity over physical spread. https://github.com/StochasticBiology/optimal-mitos/blob/main/summary-videos/several-simulations.mp4

